# Liver Viscosity Decreases Before the Onset of Fibrosis in Metabolic Dysfunction-Associated Steatohepatitis (MASH)

**DOI:** 10.1101/2025.09.24.678352

**Authors:** Yasmine Safraou, Kristin Susan Spirgath, Biru Huang, Christian Bayerl, Karolina Krehl, Anja A. Kühl, Tom Meyer, Mehrgan Shahryari, Pedro Dantas de Moraes, Jakob Jordan, Noah Jaitner, Dominik Geisel, Jörg Schnorr, Nicola Stolzenburg, Michael Mülleder, Kathrin Textoris-Taube, Iwona Wallach, Nikolaus Berndt, Heiko Tzschätzsch, Rebecca G. Wells, Jürgen Braun, Patrick Asbach, Ingolf Sack, Jing Guo

## Abstract

**Background and Aim:** Metabolic dysfunction-associated steatohepatitis (MASH) is an increasingly prevalent condition worldwide, associated with biomechanical liver changes and detectable by magnetic resonance elastography (MRE). This study explored the pathophysiological features and their biomechanical manifestations at different stages of MASH in a mouse dietary model.

**Methods:** Using MRE on a clinical 3 Tesla MRI scanner, we measured liver stiffness, viscosity, fat fraction and water diffusion in 45 male mice. These values were correlated with histopathology and proteomics analyses to further characterize the liver microstructural and metabolic changes during MASH progression.

**Results:** We found in a high-fat, low amino-acid model that early MASH was marked by fat accumulation and increasing inflammatory activity, while later stages showed a reduction in fat despite persistent inflammation. These changes in microstructure were associated with biomechanical adaptations, including a progressive decrease in hepatic viscosity and the water diffusion. Notably, viscosity was inversely correlated with lobular inflammation, cell adhesion, antioxidant activity, and metabolic adaptations such as enhanced ketone body synthesis. These findings, which precede the onset of fibrosis and tissue stiffening, show that tissue viscosity is highly sensitive to early microstructural and metabolic alterations in MASH.

**Conclusion:** Steatosis and inflammation significantly alter liver biophysical properties, particularly viscosity, in a mouse dietary model of MASH, even in the absence of fibrosis. These findings suggest that viscosity is a potential early and clinically translatable biomarker for the development and progression of MASH.

## 1. Introduction

Metabolic dysfunction-associated steatotic liver disease (MASLD), formerly known as non-alcoholic fatty liver disease (NAFLD), is the most common chronic liver disease worldwide [1, 2]. An estimated one-third of the world’s population is affected by MASLD, which parallels the increasing prevalence of obesity and type 2 diabetes mellitus [3, 4]. The recent switch in terminology from NAFLD to MASLD reflects the frequent occurrence of metabolic comorbidities [5–9]. MASLD can progress to a more severe stage known as metabolic dysfunction-associated steatohepatitis (MASH), which is characterized by active necroinflammation including lobular inflammation and hepatocyte ballooning [1, 3, 5, 10]. This sustained liver cell damage results in the development of fibrosis [1, 5, 11], which, if unchecked, may lead to cirrhosis/end-stage liver disease, hepatocellular carcinoma and extrahepatic pathology including cardiometabolic diseases [5, 12]. Liver disease progression is associated with disruptions in normal metabolic processes such as gluconeogenesis, lipid metabolism, and detoxification, further contributing to systemic complications that exacerbate the risk of cardiometabolic diseases and metabolic dysfunction-associated pathologies [5, 13].

Although liver biopsy remains the gold standard for assessing MASLD and MASH, its invasive nature and potential sampling errors have prompted the search for noninvasive imaging markers. Magnetic resonance imaging (MRI) has proven to be valuable for the quantification of the fat fraction using the Dixon method [14] and diffusion-weighted imaging (DWI) offers a complementary, sensitive approach for characterizing and differentiating steatohepatitis from fibrosis [15–20]. The biomechanical properties of the liver, including stiffness and viscosity, as quantified by magnetic resonance elastography (MRE), are highly sensitive to structural and functional tissue alterations including fibrosis [19, 21–24], steatosis [25], microstructural heterogeneities [26], cellular hypertrophy [27], altered metabolic functions [28], and fluctuations in hepatic vascular pressure [29–32] – all of which are potentially relevant for assessing disease progression in patients with MASLD and MASH [25, 26, 33]. Despite the increasing availability of quantitative and biophysical MRI techniques, there is still no established imaging biomarker to indicate the microstructural and metabolic transition from MASLD to MASH, that is from simple steatosis to the combined presence of steatosis and inflammation, before the occurrence of fibrosis. Moreover, the variability in MASLD progression among patients presents a significant challenge in linking specific pathological features with in vivo biophysical imaging properties. In contrast, a preclinical model of MASLD/MASH would facilitate effective control over disease progression while at the same time allowing establishment of histopathological and biochemical references to eventually identify predictive MASLD/MASH imaging markers for clinical use in patients.

This study employed a clinical MRI scanner equipped with an MRE modality, alongside histology and metabolic analysis based on a kinetic model of central liver metabolism [34–36], to investigate the relationship between imaging biophysical markers (including stiffness, viscosity and water diffusivity) and the histopathologic and metabolic characteristics of MASLD and MASH in a mouse dietary model. The primary objective was to identify imaging biomarkers sensitive to specific pathophysiological hallmarks of MASLD/MASH, including metabolic alterations, to enable early disease detection. Based on these data, we suggest that viscosity is a sensitive marker of pre-fibrotic disease progression.

## 2. Results

### 2.1. Histology

Steatosis, inflammation and fibrosis were initially evaluated in the mouse model. Male mice were fed a choline-deficient, L-amino acid-defined, high-fat diet (CDAHFD) beginning at 45 days of age and were studied at baseline and at multiple time points extending up to 121 days [17]. Liver histology was evaluated using two complementary approaches: an initial semiquantitative analysis with the non-alcoholic fatty liver activity score (NAS), followed by quantitative segmentation of high-resolution Hematoxylin & Eosin (H&E)-stained sections to determine percentage amounts of fat and surface densities of lobular inflammation.

Semiquantitative histology based on NAS showed that all mice fed the CDAHFD developed steatosis and inflammation as early as the first examined time point (day 12). Notably, no mouse exhibited steatosis alone, with absence of inflammation. Fibrosis was present to a lesser extent and was first observed starting from day 21.

In detail, stage 3 steatosis was observed in all mouse groups on the diet (from day 12 to day 121). Quantitative histology showed that the fat ratio fluctuated significantly over the course of the disease (*p* = 3.0e-7). A significant increase of 631 ± 1895 % in fat ratio was observed between day 0 and day 21 of dietary treatment (fat ratio _day0_ = 5.7 ± 10.6 %, fat ratio _day21_ = 41.4 ± 14.8 %, *p* = 3.5e-4), followed by a significant decrease of 18 ± 36 % (fat ratio _day121_ = 34.0 ± 1.7 %, *p* = 4.4e-4) up until day 121.

Inflammation increased with time on the diet. Portal inflammation was observed in eight of the 45 mice, whereas multifocal lobular inflammation was increasingly and consistently present in all animals from day 12 to day 121 of CDAHFD. The lobular inflammation was characterized by the presence of small foci of inflammatory cell infiltrates, most of them lymphocytes, with a smaller proportion of macrophages, and aggregates of neutrophils. The segmentation of cell infiltrate showed a consistent increase of 131 ± 77 % from day 0 to day 121 (infiltrate density _day0_ = 5.7e-4 ± 9.6e-5 µm^-2^, infiltrate density _day121_ = 13.0e-4 ± 1.8e-4 µm^-2^, *p* = 5.5e-7). The cell infiltrate density parameter strongly correlated with the semiquantitative lobular inflammation score (R = 0.76, *p* = 3.0e-9).

Fibrosis stage 1 was identified in eight mice across the cohort, whereas only two mice progressed to stage 2 fibrosis. A quantitative fibrosis score was not established as no consistent microscopic fibrosis features, such as bridging, were observed in the examined H&E-stained slides. Representative microscopic images illustrating the histological findings and quantitative histological segmentation results are shown in Figure 1. All histological data are plotted in Figure 2, with quantitative segmentation results in Figure 2(A) and semiquantitative histology scores in Figure 2(B). A comprehensive summary of all histological findings is compiled in Table 1.

**Figure 1.**
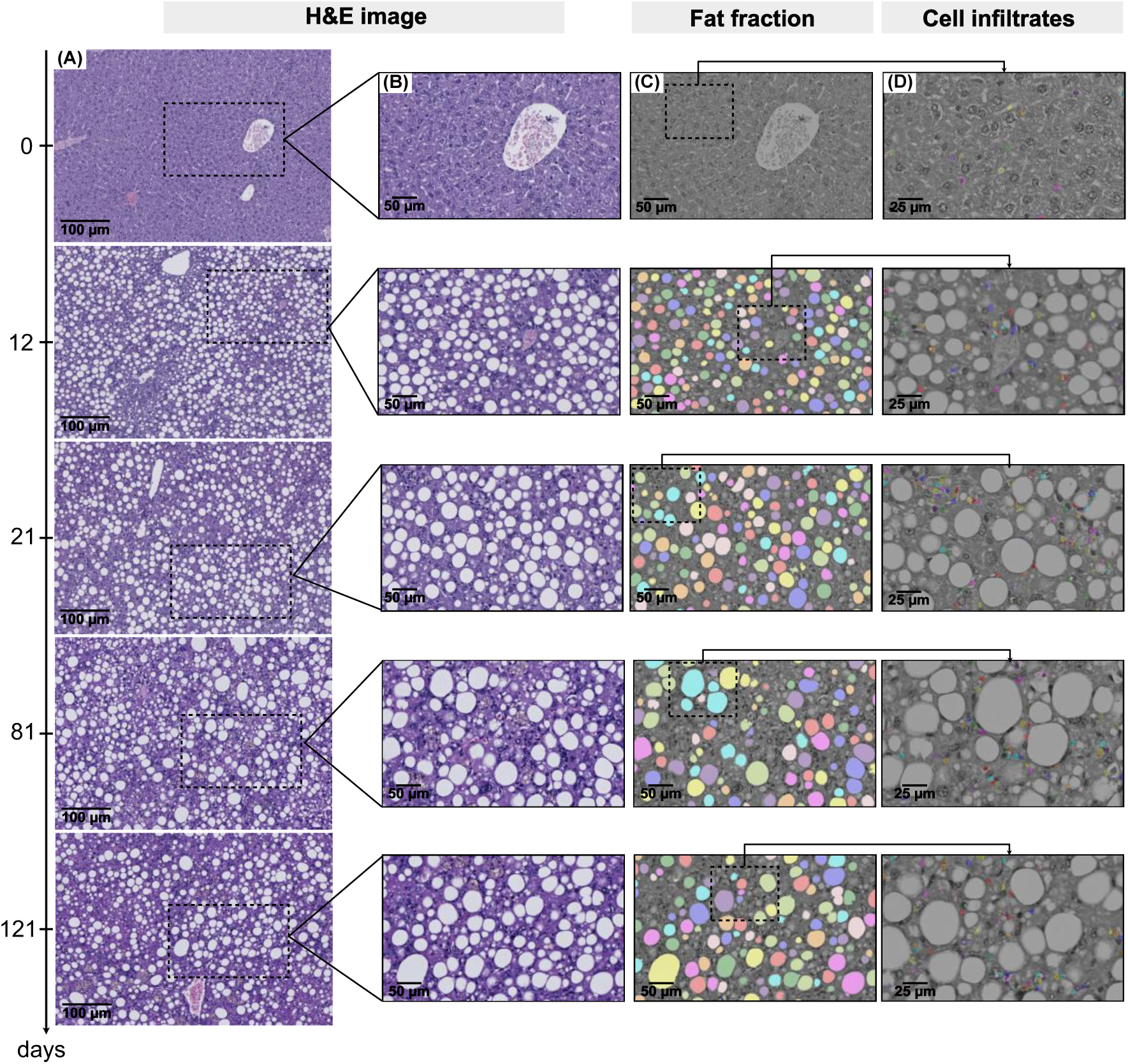
Microscopy images of Hematoxylin and Eosin (H&E)-stained liver sections across all experimental time points (0, 12, 21, 81, and 121 days) and corresponding fat and lymphocyte segmentations. (A) Representative snapshots (5000 x 5000 pixels, 228 x 228 nm² resolution) selected from high-resolution H&E images of whole liver sections. (B) Zoomed-in views showing detailed visual structures from the H&E-stained sections. (C) Grayscales image showing snapshots of the fat droplet segmentation masks; the visible colors represent clustering algorithm outputs and have no quantitative significance. (D) Further zoomed-in views from panels (B) and (C) illustrating segmented lymphocytes. As in panel (C), the visible colors correspond to clustering algorithm outputs without quantitative significance.

**Figure 2.**
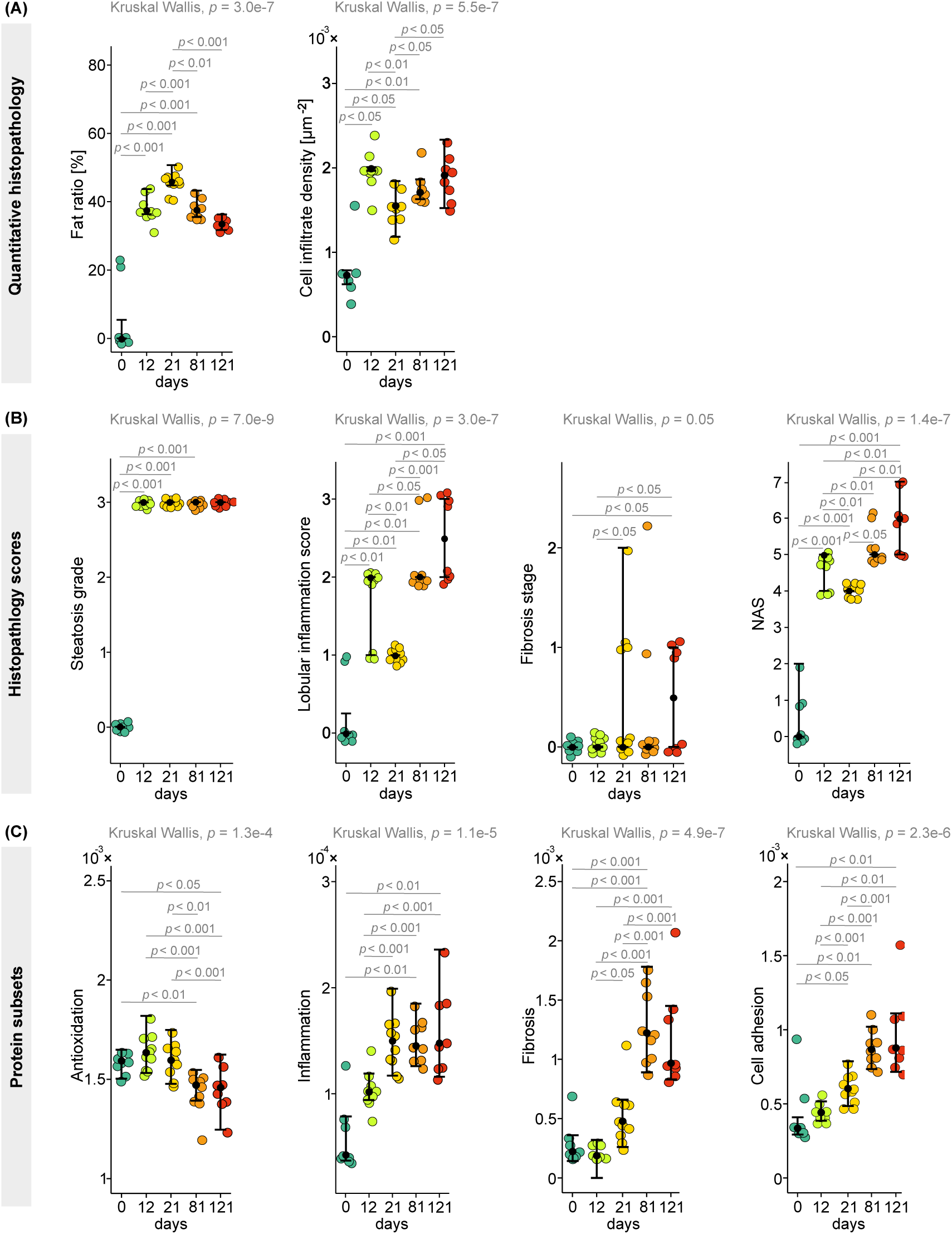
Swarm plots illustrating evolution of postmortem parameters across all experimental time points. (A) Quantitatively derived histological segmentation parameters. (B) Semiquantitative histology scores. (C) Quantitative subset ratios of all protein subsets. Differences between time points were statistically analyzed using the Kruskal-Wallis test, while post hoc pairwise comparisons were performed using the Wilcoxon-Mann-Whitney test. Mean and lower and upper quartiles are indicated for each time point. NAS: non-alcoholic fatty liver activity score.

**Table 1.** Histological assessment of liver tissue across all four time points of diet intake (day 0, day 12, day 21, day 81, and day 121 on diet). Histological scores of steatosis, lobular inflammation, fibrosis, cell ballooning and the non-alcoholic fatty liver activity score (NAS) are presented as semiquantitative grades. Quantitative scores of fat ratio (%) and cell infiltrate density (µm^-2^) are presented as means and standard deviations.

|  |  |  | Day 0 | Day 12 | Day 21 | Day 81 | Day 121 |
| --- | --- | --- | --- | --- | --- | --- | --- |
|  |  |  | N = 8 | N = 10 | N = 10 | N = 9 | N = 8 |
| Histological scores | Semiquantitative | Steatosis grade | Grade 0 (7)<br>Grade 1 (1) | Grade 3 (10) | Grade 3 (10) | Grade 3 (9) | Grade 3 (8) |
|  |  | Lobular inflammation | Grade 0 (7)<br>Grade 1 (1) | Grade 0 (3)<br>Grade 1 (7) | Grade 1 (10) | Grade 2 (7)<br>Grade 3 (2) | Grade 2 (4)<br>Grade 3 (4) |
|  |  | Fibrosis | Stage 0 (8) | Stage 0 (10) | Stage 0 (6)<br>Stage 1 (3)<br>Stage 2 (1) | Stage 0 (7)<br>Stage 1 (1)<br>Stage 2 (1) | Stage 1 (4)<br>Stage 2 (4) |
|  |  | Cell ballooning | Score 0 (7)<br>Score 1 (8) | Score 0 (10) | Score 0 (10) | Score 0 (8)<br>Score 1 (1) | Score 0 (7)<br>Score 1 (1) |
|  |  | NAS | Score 0 (5)<br>Score 1 (2)<br>Score 2 (1) | Score 4 (3)<br>Score 5 (7) | Score 4 (10) | Score 5 (7)<br>Score 6 (1)<br>Score 7 (1) | Score 5 (3)<br>Score 6 (3)<br>Score 7 (3) |
|  | Quantitative | Fat ratio (%) | 5.7 ± 10.4 | 38.6 ± 3.9 | 46.4 ± 3.2 | 38.3 ± 2.8 | 34.0 ± 1.7 |
| | | Cell infiltrate density ($\mu\text{m}^{-2}$) | 5.6e-4 ± 9.6e-5 | 7.5e-4 ± 6.2e-5 | 8.0e-4 ± 6.4e-5 | 11.4 e-4 ± 1.7e-4 | 13.03 ± 1.8e-4 |

### 2.2. Imaging-derived biophysical properties of the liver

Animals were imaged with a clinical MRI scanner at each timepoint of the study, prior to post-mortem histological analysis. Figure 3(A) shows representative examples of anatomical T2-weighted (T2w) images and maps of shear wave speed (*SWS*), penetration rate (*PR*), hepatic fat fraction (*HFF*), and apparent diffusion coefficient (*ADC*). *SWS* maps serve as a surrogate for liver stiffness, while *PR* maps reflect inverse viscosity. *ADC* maps surrogate hepatic water diffusion.

**Figure 3.**
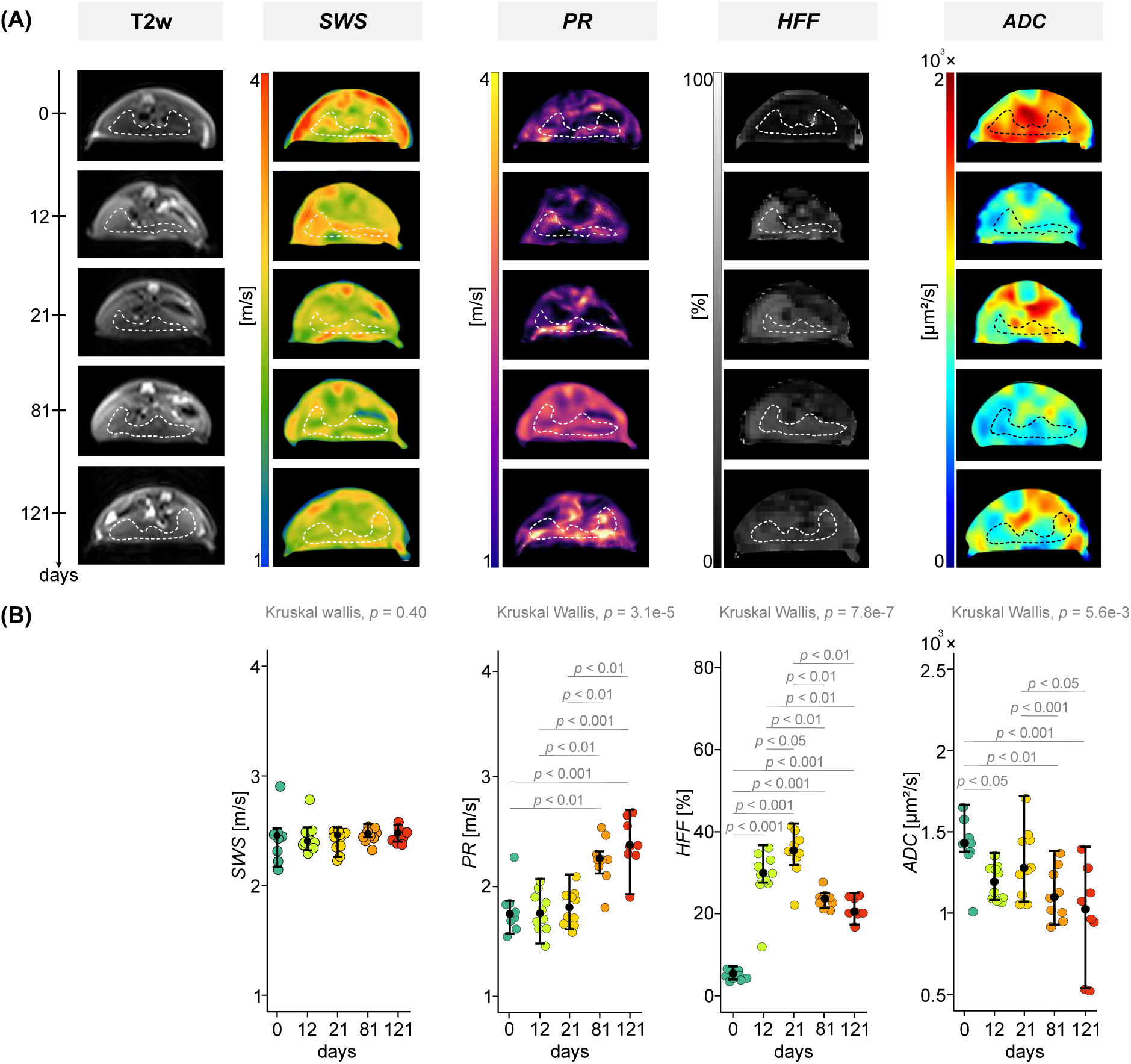
Abdominal imaging results. (A) Examples of T2-weighted (T2w) images, *SWS* maps, *PR* maps, *HFF* maps, and *ADC* maps acquired at all five experimental time points in a total of 45 mice: baseline (n=8), and after 12 (n = 10), 21 (n = 10), 81 (n = 9), and 121 days (n = 8) on diet. The liver is demarcated by a dotted white or black line. (B) Swarm plots of the evolution of imaging parameters (*SWS*, *PR*, *HFF* and *ADC*) over all time points show means with error bars indicating lower and upper quartiles. Differences between time points were statistically analyzed using the Kruskal-Wallis test. Post hoc pairwise comparisons were performed using the Wilcoxon-Mann-Whitney test. *SWS*: shear wave speed, *PR*: penetration rate, *HFF*: hepatic fat fraction, *ADC*: apparent diffusion coefficient.

The maps demonstrate an increase in *PR* and a decrease in *ADC* throughout the course of disease progression. *HFF* showed a steady increase from baseline (day 0, control group) to day 21, followed by a gradual decline until day 121. No changes in *SWS* maps were visible.

Consistent with the visual observations, quantitative imaging values averaged from liver regions of interest (ROIs) exhibited similar trends:

Hepatic *PR* (inverse viscosity) values increased by 34 ± 22 % from day 0 to day 121 in the cohort as a whole (*PR* _day0_ = 1.81 ± 0.22 ms, *PR* _day121_ = 2.41 ± 0.24 ms, *p* = 3.1e-5).

*HFF* fluctuated significantly from day 0 to day 121 (*p* = 7.8e-7). Between day 0 and day 21, *HFF* showed a significant increase of 553 ± 200 % (*HFF* _day0_ = 5.4 ± 1.0 %, *HFF* _day21_ = 35.5 ± 5.4 %, *p* = 4.5e-4), followed by a decrease of 39 ± 16 % from day 21 to day 121 (*HFF* _day121_ = 21.9 ± 2.7 %, *p* = 3.2e-4).

For *ADC* (water diffusion), a consistent decrease was observed over time with a significant reduction of 31 ± 21% between day 0 and day 121 (*ADC* _day0_ = 1427.0 ± 191 µm²/s, *ADC* _day121_ = 995 ± 317 µm²/s, *p* = 5.6e-3).

There were no significant differences in *SWS* (stiffness) (*p* = 0.40). The group analysis results for all imaging parameters are plotted in Figure 2(B) and compiled in Table 2 below.

**Table 2.**
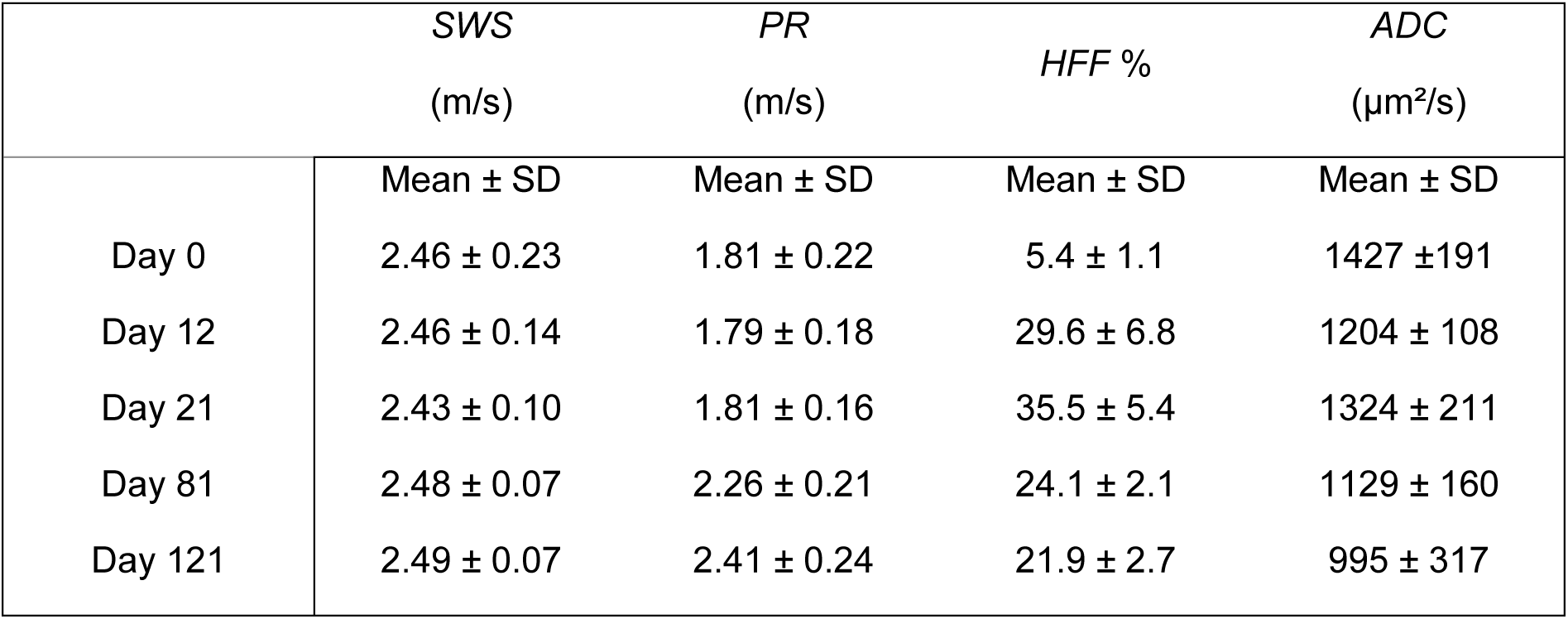
Group mean values and standard deviations of imaging parameters - shear wave speed (*SWS*), penetration rate (*PR*), hepatic fat fraction (*HFF*), and apparent diffusion coefficient (*ADC*) - measured across all mice groups at different time points: control (day 0), day 12, day 21, day 81, and day 121. SD stands for standard deviation.

### 2.3. Proteomics analyses

#### Metabolic capacities

Proteomics analyses were performed on snap-frozen liver specimens. The metabolic signatures were derived using HEPATOKIN 1 [34], a kinetic model that allows a comprehensive evaluation of the key pathways involved in carbohydrate, lipid, amino acid, and energy metabolism in hepatocytes. The model quantifies physiological metabolic functions as maximal fluxes across a range of conditions, spanning from fasted (low glucose, high FA, low insulin, and high glucagon) to fed states (high glucose, low FA, high insulin, and low glucagon). Metabolic functions describing fluxes are reported in µmol/g/h, while functions related to metabolite uptake were expressed in mM, indicating extracellular substrate availability. In total, twelve metabolic functions showed significant differences across the samples analyzed in our study.

An increase in triacylglycerol (TAG) content was observed from day 0 to day 21 (TAG content _day0_ = 16.3 ± 6.3 mMol, TAG content _day21_ = 35.8 ± 5.5 mMol, *p* = 4.6e-5), followed by a subsequent decrease between day 21 and day 121 (TAG content _day121_ = 16.8 ± 8.6 mMol, *p* = 4.6e-5). This change in TAG content indicates an adaptation to the high-fat diet and corresponds to alterations in hepatic fat content.

A significant increase in ketone body production was observed on the diet (ketone bodies _day0_ = 30.2 ± 12.3 µmol/g/h, ketone bodies _day121_ = 38.0 ± 6.4 µmol/g/h, *p* = 8.7e-4). Among the quantified ketone bodies, beta-hydroxybutyrate (BHBs) increased significantly (BHBs _day0_ = 8.0 ± 5.3 µmol/g/h, BHBs _day121_ = 23.0 ± 7.5 µmol/g/h, *p* = 3.5e-5). Urea production increased significantly from day 0 to day 12 (urea production _day0_ = 44.2 ± 17.6 µmol/g/h, urea production _day12_ = 58.3 ± 2.8 µmol/g/h, *p* = 6.0e-3) although it then decreased between day 12 and day 121 (urea production _day121_ = 27.8 ± 14.0 µmol/g/h, *p* = 9.1e-5). These findings indicate a profound metabolic reprogramming and a potential impairment in liver function simultaneously with the progression of MASLD during dietary treatment. A detailed group analysis of significantly altered maximal metabolic functions is presented in Table 3(A). An extensive description of the metabolic model is provided in the section “Metabolic profiling” of the Supplemental material (See Figure S1 and Table S1). Boxplots illustrating these functions are shown in Figure S2 of the Supplemental material.

**Table 3.** Summary of results of proteomics analysis. Results are provided as mean and standard deviation (SD) for (A) metabolic functions and (B) protein subsets representing antioxidation, inflammation, fibrosis, and cell adhesion for baseline and all four time points of diet intake. Differences between time points were statistically analyzed using the Kruskal-Wallis test. FA stands for fatty acid, TAG for triacylglycerol, VLDL for very-low-density lipoprotein, BHB for beta-hydroxybutyrate, and ACAC for acetoacetate.

| (A) Metabolic functions |  |  |  |  |  |  |
| --- | --- | --- | --- | --- | --- | --- |
|  | Day 0 | Day 12 | Day 21 | Day 81 | Day 121 | *p* |
|  | Mean ± SD | Mean ± SD | Mean ± SD | Mean ± SD | Mean ± SD |  |
| FA uptake<br>(μmol/g/h) | 30.9 ± 16.0 | 39.8 ± 14.6 | 37.7 ± 15.1 | 36.3 ± 12.6 | 31.5 ± 9.8 | 0.035 |
| TAG synthesis<br>(μmol/g/h) | 13.9 ± 5.1 | 14.5 ± 0.7 | 14.7 ± 3.2 | 11.2 ± 3.2 | 8.0 ± 4.0 | 1.7e-4 |
| TAG content<br>(mMol) | 16.3 ± 6.3 | 34.5 ± 6.6 | 35.8 ± 5.5 | 23.5 ± 8.1 | 16.8 ± 8.6 | 5.3e-6 |
| VLDL export<br>(μmol/g/h) | 4.4 ± 1.8 | 6.1 ± 0.5 | 5.9 ± 1.0 | 5.0 ± 1.5 | 3.5 ± 1.7 | 1.7e-3 |
| Ketone body<br>production<br>(μmol/g/h) | 30.2 ± 12.3 | 33.0 ± 6.7 | 27.5 ± 5.7 | 39.0 ± 3.8 | 38.0 ± 6.4 | 8.7e-4 |
| BHB production<br>(μmol/g/h) | 8.0 ± 5.3 | 7.7 ± 2.5 | 7.8 ± 1.8 | 15.8 ± 6.3 | 23.0 ± 7.5 | 3.5e-5 |
| ACAC production<br>(μmol/g/h) | 19.5 ± 9.2 | 24.6 ± 2.3 | 20.0 ± 4.3 | 26.2 ± 3.5 | 18.0 ± 7.4 | 9.3e-3 |
| Ammonia uptake<br>(mMol) | 128.7 ±<br>23.6 | 153.0 ± 5.6 | 162.42 ±<br>7.9 | 144.0 ± 19.0 | 137.6 ±<br>39.4 | 2.3e-3 |
| Urea production<br>(μmol/g/h) | 44.2 ± 17.6 | 58.3 ± 2.8 | 53.9 ± 7.9 | 39.3 ± 16.3 | 27.8 ± 14.0 | 2.7e-5 |
| Galactose uptake<br>(mMol) | 20.5 ± 5.8 | 20.1 ± 2.5 | 23.3 ± 3.1 | 22.7 ± 8.5 | 12.0 ± 8.3 | 0.02 |
| Fructose uptake<br>(μmol/g/h) | 367.2 ±<br>46.6 | 258.0 ±<br>22.0 | 217.4 ±<br>22.6 | 233.2 ±<br>53.0 | 230.0 ±<br>16.6 | 2.0e-5 |
| Gluconeogenesis<br>(μmol/g/h) | 64.8 ± 27.9 | 87.2 ± 9.4 | 82.6 ± 6.7 | 58.2 ± 29.6 | 27.6 ± 17.2 | 6.7e-5 |
| (B) Protein subsets |  |  |  |  |  |  |
|  | day0 | day12 | day21 | day81 | day121 | *p* |
|  | Mean ± SD | Mean ± SD | Mean ± SD | Mean ± SD | Mean ± SD |  |
| Antioxidation | 1.6e-2 ±<br>5.2e-4 | 1.7e-2 ±<br>9.0e-4 | 1.6e-2 ±<br>8.5e-4 | 1.5e-2 ±<br>1.0e-3 | 1.5e-3 ±<br>1.2e-3 | 1.5e-4 |
| Inflammation | 6.0e-5 ±<br>3.2e-5 | 1.1e-4 ±<br>1.8e-5 | 1.5e-4 ±<br>2.6e-5 | 1.5e-4 ±<br>1.9e-5 | 1.6e-4 ±<br>4.1e-5 | 1.0e-5 |
| Fibrosis | 3.0e-4 ±<br>1.8e-4 | 1.7e-4 ±<br>1.3e-4 | 5.5e-4 ±<br>2.5e-4 | 1.3e-3 ±<br>3.2e-4 | 1.2e-3 ±<br>4.3e-4 | 3.6e-7 |
| Cell adhesion | 4.3e-4 ±<br>2.3e-4 | 4.5e-4 ±<br>5.8e-5 | 6.2e-4 ±<br>9.9e-5 | 8.9e-4 ±<br>1.3e-4 | 9.8e-4 ±<br>2.8e-4 | 2.3e-6 |

#### Evaluation of protein subsets for inflammation, antioxidant capacity, fibrosis, and cell adhesion

Significant differences over the course of MASH were also observed in proteomic subsets reflecting antioxidation, inflammation, fibrosis, and cell adhesion, listed in Table S2. Increases from day 0 to day 121 were observed in proteins related to inflammation (inflammation _day0_ = 6.0e-5 ± 3.2e-5; inflammation _day121_ = 1.6e-4 ± 4.1e-5, *p* = 1.0e-5), fibrosis (fibrosis _day0_ = 3.0e-3 ± 1.8e-3; fibrosis _day121_ = 1.2e-3 ± 4.3e-4, *p* = 3.6e-7) and cell adhesion (cell adhesion _day0_ = 4.3e-4 ± 2.3e-4; cell adhesion _day121_ = 9.8e-4 ± 2.8e-4, *p* = 2.3e-6). However, the antioxidation protein subset decreased over this time (antioxidation _day0_ = 0.016 ± 5.2e-4, antioxidation _day121_ = 1.5e-3 ± 1.2e-3, *p* = 1.5e-4), indicating higher oxidative stress, likely of hepatocytes. All results of protein subset analysis are plotted in Figure 2(C) and compiled in Table 3(B).

### 2.4. Correlations between imaging parameters, histology, and proteomics

Imaging markers correlated with both semiquantitative and quantitative histological parameters. *PR* showed a strong correlation with lobular inflammation (R = 0.55, *p* = 8.2e-5) and NAS (R = 0.55, *p =* 9.1e-5). In addition, *PR* exhibited a strong correlation with infiltrate density (R = 0.69, *p* = 2.4e-7). From day 21 to day 121, *PR* showed a strong positive correlation with infiltrate density (R = 0.6, *p* = 9.0e-4) and a strong negative correlation with the fat ratio (R = −0.82, *p* = 1.2e-7).

No correlation was observed between *SWS* and any histological scoring parameters. Strong correlations were observed between *ADC* and both the lobular inflammation score (R = −0.52, *p =* 2.5e-5) and the NAS (R = −0.59, *p* = 2.9e-5). Moderate correlation was obtained between *ADC* and the steatosis grade (R = −0.45, *p* = 2.1e-3) as well as infiltrate density (R = −0.39, *p* = 9.0e-3).

*HFF* measured by MRI strongly correlated with the steatosis grade (R = 0.66, *p* = 7.3e-7). Furthermore, *HFF* strongly correlated with the quantitative fat ratio (R = 0.78, *p* = 5.2e-10).

Correlations between imaging parameters and the four protein subsets identified by proteomics analyses (antioxidation, inflammation, fibrosis, and cell adhesion) were identified exclusively for *PR* and *ADC*. Most notably, there were strong correlations between *PR* and protein subsets of cell adhesion (R = 0.55, *p* = 7.6e-5), fibrosis (R = 0.58, *p* = 2.5e-5), and antioxidation (R = −0.52, *p* = 3.0e-4). Additionally, a moderate correlation was observed between *PR* and the inflammation subset (R = 0.40, *p* = 7.5e-3).

Correlations between imaging parameters and maximum metabolic capacities were predominantly observed for *PR* and *HFF*. *PR* correlated strongly with the maximal capacity for ketone body production (R = 0.56, *p* = 6.7e-5) and for BHB production (R = 0.66, *p* = 9.4e-7).

Weak correlations were found between *PR* and the maximal capacity for TAG synthesis (R = −0.43, *p* = 2.8e-3), urea production (R = −0.47, *p* = 1.2e-3), and gluconeogenesis (R = −0.46, *p* = 1.7e-3). There was a strong correlation between MRI-based *HFF* and both maximal TAG content (R = 0.78, *p* = 3.3e-10) and maximal capacity for ammonia uptake (R = 0.64, *p* = 1.9e-6). Moderate correlations were found between *HFF* and maximal capacities for FA uptake (R = 0.35, *p* = 0.02), very-low density lipoprotein (VLDL) export (R = 0.45, *p* = 1.6e-3), urea production (R = 0.46, *p* = 1.3e-3), gluconeogenesis (R = 0.40, *p* = 6.5e-3), and fructose uptake (R = −0.47, *p* = 1.1e-3). Graphs for the most relevant correlations are presented in Figure 4, while a comprehensive overview of correlations between imaging biomarkers and post-mortem tissue parameters is provided in Table 4.

**Figure 4.**
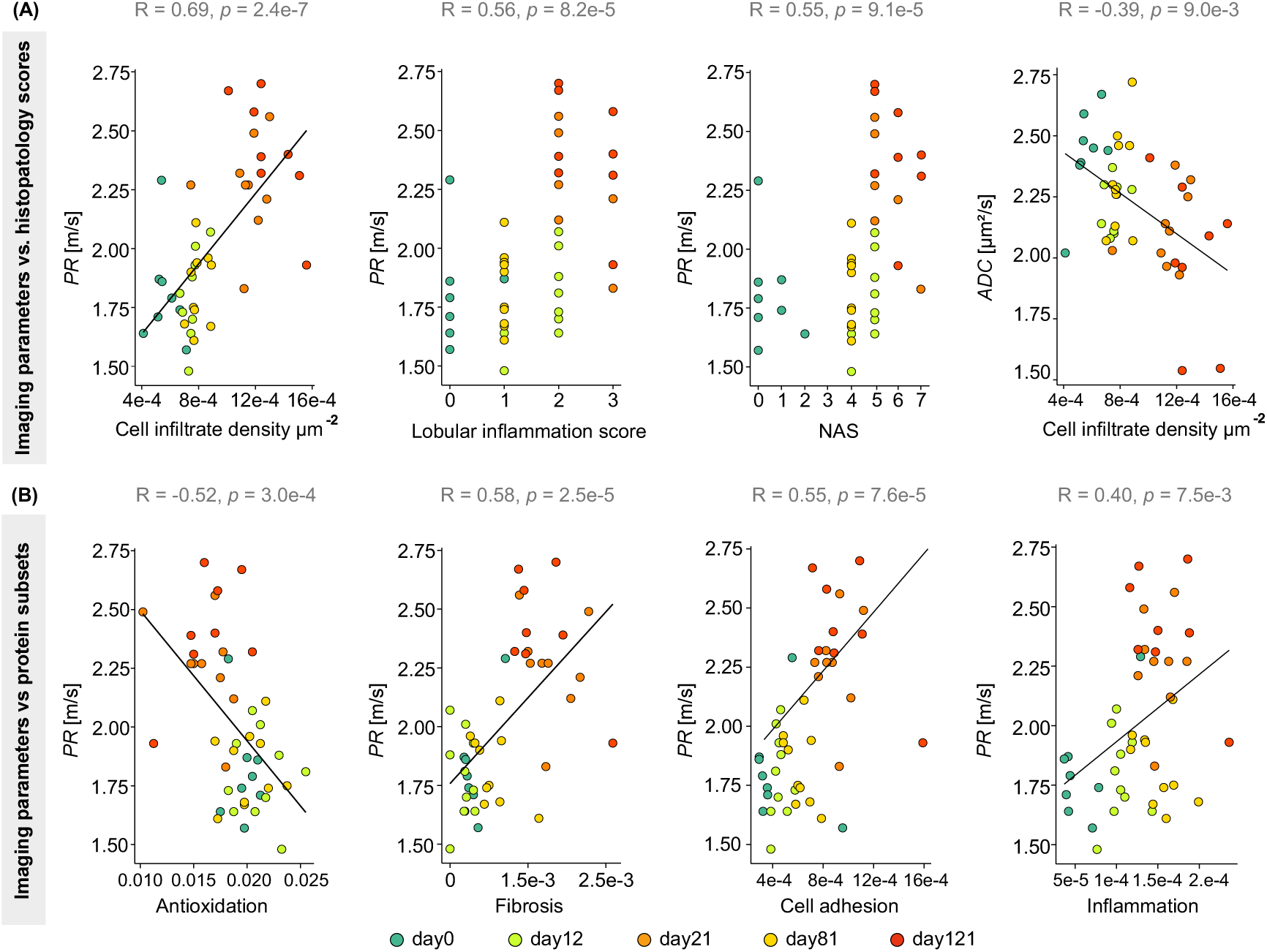
Relevant correlations between imaging parameters and postmortem tissue parameters. Spearman correlation plots are presented, showing imaging parameters on the y-axis and quantitative or semiquantitative histopathology scores on the x-axis (A) or protein subsets associated with several functions (B).

**Table 4.** Results of Spearman correlation analysis for associations between imaging parameters and postmortem tissue parameters.

|  |  | *SWS* | *PR* | *HFF* | *ADC* |
| --- | --- | --- | --- | --- | --- |
| Histology | Lobular inflammation | $p = 0.29$ | $R = 0.56$<br>$p = 8.2e-5$ | $p = 0.63$ | $R = -0.52$<br>$p = 2.5e-5$ |
| | Steatosis grade | $p = 0.60$ | $R = 0.30$<br>$p = 0.05$ | $R = 0.66$<br>$p = 7.3e-7$ | $R = -0.45$<br>$p = 2.1e-3$ |
| | Fibrosis | $p = 0.60$ | $p = 0.22$ | $p = 0.84$ | $p = 0.17$ |
| | NAS | $p = 0.31$ | $R = 0.55$<br>$p = 9.1e-5$ | $p = 0.47$ | $R = -0.58$<br>$p = 2.9e-5$ |
| | Fat ratio (%) | $p = 0.31$ | $p = 0.31$ | $R = 0.78$<br>$p = 5.2e-10$ | $p = 0.52$ |
| | Cell infiltrate density<br>[$\mu m^{-2}$] | $p = 0.40$ | $R = 0.69$<br>$p = 2.4e-7$ | $p = 0.44$ | $R = -0.39$<br>$p = 9.0e-3$ |
| Protein subsets | Cell adhesion | $p = 0.09$ | $R = 0.55$<br>$p = 7.6e-5$ | $p = 0.99$ | $R = -0.35$<br>$p = 0.02$ |
| | Inflammation | $p = 0.72$ | $R = 0.40$<br>$p = 7.5e-3$ | $R = 0.32$<br>$p = 0.04$ | $R = -0.37$<br>$p = 0.02$ |
| | Antioxidation | $p = 0.95$ | $R = -0.52$<br>$p = 0.0003$ | $p = 0.21$ | $R = 0.31$<br>$p = 0.04$ |
| | Fibrosis | $p = 0.33$ | $R = 0.58$<br>$p = 2.5e-5$ | $p = 0.95$ | $R = -0.31$<br>$p = 0.04$ |
| Maximum metabolic functions | FA uptake [$\mu\text{mol/g/h}$] | $p = 0.35$ | $p = 0.81$ | $R = 0.35$<br>$p = 0.02$ | $p = 0.65$ |
| | TAG synthesis<br>[$\mu\text{mol/g/h}$] | $R = -0.34$<br>$p = 0.03$ | $R = -0.43$<br>$p = 2.8\text{e-}3$ | $p = 0.26$ | $R = 0.40$<br>$p = 6.5\text{e-}3$ |
| | TAG content [mMol] | $p = 0.14$ | $p = 0.19$ | $R = 0.78$<br>$p = 3.3\text{e-}10$ | $p = 0.12$ |
| | VLDL export<br>[$\mu\text{mol/g/h}$] | $p = 0.17$ | $p = 0.19$ | $R = 0.45$<br>$p = 1.6\text{e-}3$ | $p = 0.48$ |
| | Ketone body<br>production [$\mu\text{mol/g/h}$] | $p = 0.49$ | $R = 0.56$<br>$p = 6.7\text{e-}5$ | $p = 0.30$ | $R = -0.31$<br>$p = 0.04$ |
| | BHB production<br>[$\mu\text{mol/g/h}$] | $p = 0.49$ | $R = 0.66$<br>$p = 9.4\text{e-}7$ | $p = 0.18$ | $p = 0.12$ |
| | ACAC production<br>[$\mu\text{mol/g/h}$] | $p = 0.07$ | $p = 0.94$ | $p = 0.93$ | $p = 0.43$ |
| | Ammonia uptake<br>[mMol] | $p = 0.15$ | $p = 0.41$ | $R = 0.64$<br>$p = 1.9\text{e-}6$ | $p = 0.82$ |
| | Urea production<br>[$\mu\text{mol/g/h}$] | $R = -0.30$<br>$p = 0.05$ | $R = -0.47$<br>$p = 1.2\text{e-}3$ | $R = 0.46$<br>$p = 1.3\text{e-}3$ | $p = 0.28$ |
| | Galactose uptake<br>[mMol] | $p = 0.11$ | $p = 0.22$ | $p = 0.16$ | $p = 0.65$ |
| | Fructose uptake<br>[μmol/g/h] | $p = 0.45$ | $p = 0.45$ | $R = -0.47$<br>$p = 1.1\text{e-}3$ | $p = 0.12$ |
| | Gluconeogenesis<br>[μmol/g/h] | $p = 0.13$ | $R = -0.46$<br>$p = 1.7\text{e-}3$ | $R = 0.40$<br>$p = 6.5\text{e-}3$ | $p = 0.33$ |

## 3. Discussion

In this study, we show that disease progression in a mouse model of MASH strongly correlates with in vivo biophysical parameters including water diffusion and viscosity, and with metabolic factors including antioxidant activity and ketone body synthesis. Notably, changes in biophysical parameters as measured by MRI/MRE occurred as inflammation increased, but before the development of fibrosis. These findings suggest that MRI/MRE-derived changes in viscosity and water diffusivity should be evaluated as a marker of MASLD patients at risk of progression.

### Dynamic shifts in hepatic fat and inflammation drive two-stage MASH progression and metabolic reprogramming

Histological findings showed that MASH progression in our CDAHFD-fed mice did not follow a sequential progression from steatosis to inflammation. Instead, it involved a rapid and co-occurrent onset of steatosis and inflammation evident as early as day 12. This accelerated inflammatory cascade is likely driven by choline and methionine deficiency known to impair triglyceride export to the bloodstream [4, 8]. Correspondingly, we observed intrahepatocellular accumulation of free FAs and increased oxidative stress, both of which promote the release of proinflammatory cytokines and the recruitment of inflammatory cells [4, 5, 8, 37, 38].

Another key feature of our model is the biphasic variation of in vivo *HFF*, which increased notably from day 0 to day 21, then significantly declined from day 21 to day 121. This pattern supports a two-stage development of hepatic fat metabolism characterized by storage and decomposition. A similar trend was observed in ex vivo fat ratio data from quantitative histology and was further corroborated by our metabolic analysis which showed fluctuations in FA uptake, TAG synthesis, VLDL export, and ketone body production over the entire disease course. [38–41].

Specifically, FA uptake decreased significantly as early as day 21, suggesting a ‘saturated’ fat intake despite a continuous CDAHFD. This reduction may reflect a compromised hepatic function in the context of oxidative stress, hepatocytic apoptosis and subsequent clearance of fat-laden cells [42–45]. After day 21, a transition in the FA partition towards ketone body production, particularly BHBs, occurred at the expense of VLDL-TAG synthesis. This metabolic remodeling may result from higher inflammation and compromised liver function, as further evidenced by the decrease in urea production [39, 41, 42, 46–50].

It is important to note that our metabolic analysis focused on evaluating hepatic metabolic capacities rather than real-time metabolic fluxes. This standardized strategy enabled a quantitative assessment of liver metabolic functions while minimizing susceptibility to short-term metabolic fluctuations.

Collectively, these findings suggest that MASH progression in response to CDAHFD involves an inflammation-driven microstructural and metabolic reprogramming, promoting a shift in liver viscoelastic properties towards soft-solid behavior.

### Correlation between MR-based biophysical parameters and MASH features

Hepatic water diffusion, as reflected by the *ADC*, decreased as MASH developed and progressed. A negative correlation was observed between *ADC* and both steatosis grade and lobular inflammation score. The observed reduction in *ADC* in more advanced (but still non-fibrotic) MASH may be associated with the accumulation of inflammatory cell infiltrates and intracellular fat droplets [51, 52]. Fibrosis was observed in only a few mice. As expected, liver stiffness showed only minor fluctuations, indicated by unchanged *SWS* values and only minor increases in the storage modulus *G*’ (Figure S3(A)). Nevertheless, proteomics analysis revealed an upregulation of fibrosis-associated proteins, suggesting that microscopic collagen accumulation may have occurred, albeit with minimal crosslinking. As reported in the literature, increased liver stiffness is typically associated with collagen crosslinking, the presence of isolated thick collagen fibers, periportal fibrous spikes, fibrosis in the portal tract, and an increased number of fibrous septa [15, 19, 21, 23, 25, 26, 53–59]. These hallmarks were not present at any stage in our model, including in mice with stage 2 fibrosis. This likely explains why *SWS* values in these animals remained comparable to their non-fibrotic counterparts.

The marked reduction in viscosity, as indicated by an elevated *PR*, trend towards lower *G*’’ (Figure S3(B)), and reduced loss angle (Figure S3(D)), was perhaps our most surprising finding. The reduction in viscosity was strongly correlated with the liver’s capacity to produce ketone bodies and BHBs, the latter a form of primary ketone body that is synthetized and released by the liver for use as an energy substrate by other organs [50]. Although the relationship between BHBs and MASH has not been well studied, two papers report that, consistent with our findings, BHB production was associated with the activation of an anti-inflammatory response [50, 60]. The sensitivity of tissue viscosity to inflammation was also consistent with the correlation between *PR* and protein subsets associated with inflammation, antioxidation, and cell adhesion. In particular, reduced antioxidant activity and upregulated cell adhesion have been associated with greater inflammatory activity [61–63]. The increase in cell adhesion together with the recruitment and retention of immune cells may result in a more compact extracellular matrix and explain why we observed a reduction in liver viscosity.

We observed a weak positive correlation between viscosity and the histological grade of steatosis. This is in contrast to previous studies reporting increased liver viscosity due to accumulation of fat [25, 64]. We found, however, that there was a robust correlation between *PR* and inflammation and that inflammation and steatosis were always found in the same liver. Based on these findings, we hypothesized that the combination of these two factors affects tissue viscosity. To test this hypothesis, the disease course was divided into two phases: phase one (day 0 to day 21) and phase two (day 21 to day 121). While inflammatory activity increased in both phases, steatosis increased in phase one and decreased in phase two. Separate correlation analysis for the two phases showed no correlation between *PR* and inflammation or steatosis in phase one. However, in phase two, *PR* was significantly correlated with both inflammation (positively) and steatosis (negatively). These findings suggest that inflammation and steatosis may have opposing effects on viscosity – an increase in inflammation results in a decrease in hepatic viscosity, while an increase in fat content leads to increased hepatic viscosity. Thus, in phase one, the simultaneous increase in both inflammation and steatosis masked their net effects on viscosity, resulting in the absence of a correlation. In phase two, higher inflammation and lower fat content would have both led to decreased viscosity, leading to a strong correlation between the parameters. These findings suggest that, in general, inflammation has a significant and dominating influence on tissue viscosity. Salameh *et al.* observed, in a steatohepatitis rat model, that viscosity increased when isolated steatosis was present [15], which is in agreement with our results. Using a MASLD/MASH mouse model, Yin *et al.* reported an increasing damping ratio with increasing steatosis and fibrosis [53]. Based on their findings, Yin *et al.* suggested that the damping ratio, a parameter similar to the loss angle, could be used to detect inflammation in early MASLD. However, due to overlapping disease patterns, the influence of pathological features on MRE parameters is multifactorial. A recent study of Khalfallah *et al.* performed in MASLD/MASH mouse models identified weak positive correlations between viscosity based on *G*’’ and both inflammation and steatosis [65], which is in disagreement with our findings. This discrepancy could be due to the inclusion of two MASLD/MASH models and the pooling of time points across disease stages by Khalfallah *et al.* [65].

### Study Limitations

Our study has several limitations. Due to small sample sizes at the different time points, further validation is required in more animals and, ultimately, in patients using similar MRE scan protocols. Additionally, while confounding effects of fibrosis could be ruled out in our model, the concomitant development of steatosis and inflammation still presented a challenge since these features appeared to have divergent effects on the liver biophysical properties, although we were able to partially disentangle them.

## 4. Conclusion

In conclusion, the high-fat diet mouse model of MASH employed in this study enable us to investigate the effects of steatosis and inflammation on liver biophysical properties over an extended time period without the possible confounding effects of fibrosis. Our findings indicate that steatosis and inflammation, although they have opposing effects, both influence viscosity rather than stiffness. While steatosis increases viscosity, inflammation-induced upregulation of cell adhesion and immune cell infiltration causes a reduction in tissue viscosity. Nonetheless, MRE, when combined with diffusion-weighted MRI, provides a set of imaging markers for the noninvasive assessment of cell density and cellular metabolic changes, including increased ketone body synthesis. The combined use of in vivo biophysical MRI including MRE and ex vivo histology with proteomics analysis allowed us to define the progression of MASH from the structure-function perspective. The use of a clinical MRI scanner for our experimental study is conductive to clinical translation of the imaging biomarkers identified here. Our work suggests that clinical MRI/MRE studies aimed at early detection of MASLD progression, pre-fibrosis, are warranted.

## 5. Methods

### 5.1. Animal model

Animal experiments were approved by the local authorities (Landesamt für Gesundheit und Soziales Berlin, Reg. No. G 0207/20). Animal maintenance complied with national and institutional guidelines. The animals were housed in standard cages with an enriched environment (bedding and nesting material, shelters, and plastic tubes).

Forty-five C57BL/6NCrl (Charles River Laboratories, Sulzfeld, Germany) male mice (age = 45.9 ± 6.5 days, weight = 20.6 ± 2.3 g) were fed a CDAHFD (Diet reference A02082002BR, RESEARCH DIETS, New Brunswick, USA) [66]. They were divided into four groups based on different lengths of time on this diet: 12 days, n = 10; 21 days, n = 10; 81 days n = 9; 121 days, n = 8. A group of eight mice fed a normal chow diet was established as a control group (day0, n = 8). During MR imaging, the mice were anesthetized with vaporized isoflurane (1 kg/4 L/min) through a face mask attached to the bite bar. The mice were placed on a fixation table. Their eyes were hydrated (Pan-Opthal Gey Eye, Dr. Winzer Pharma GmbH, Berlin), and body temperature was maintained by placing a warm water-filled glove around their extremities.

### 5.2. Mp-MRI and MRE data acquisition

Mice were scanned in a 3-Tesla MRI scanner (Magnetom Lumina, Siemens, Germany) using a 2-receive-channel surface array coil (mouse heart surface coil array, RAPID Biomedical, Rimpar, Germany). The mp-MRI protocol included clinical T2w, DWI, and a 2-point Dixon method. All MR images were acquired in transverse slice orientation and covered the entire liver volume.

#### 5.2.1. T2w imaging

Twenty turbo-spin-echo (TSE) T2w slices with an in-plane resolution of 0.25 × 0.25 mm² and a slice thickness of 1.2 mm were acquired for anatomical references. Further imaging parameters were: echo time (TE), 2.37 ms; repetition time (TR), 15.0 ms; field of view (FoV), 150 × 150 mm², no slice gap. Total acquisition time was 6 min.

#### 5.2.2. Two-point Dixon imaging

Hepatic fat was quantified using the 2-point Dixon method in 20 slices acquired with no slice gap. Acquisition parameters were as follows: pixel size, 1.0 × 1.0 mm^2^; slice thickness, 1.0 mm; TE, 2.46 ms (out-of-phase (OP)), 3.69 (in-phase (IP)); TR, 5.6; ms; FoV, 76 × 128 mm². Total acquisition time was 20 sec.

#### 5.2.3. Diffusion-weighted imaging

DWI with a multi-shot readout-segmented echo-planar imaging (RESOLVE) sequence was performed for quantifying water diffusion. Ten slices were acquired at b-values of 0, 50, 400, and 800 s/mm². These b-values were selected based on clinical and preclinical imaging protocols as recommended by Safraou *et al.* [29]. Two echo times were used: TE _imaging_ = 47 ms for image acquisition, and TE _navigator_ = 61 ms for k-space navigation. TR was set at 1890 ms. In-plane resolution was 1.0 × 1.0 mm³ and slice thickness was 2.3 mm. FoV was 64 × 164 mm². Parallel imaging was performed with a GRAPPA factor of 2. The averages of the diffusion coefficients for each b-value were 5 for 0, 50, and 400 s/mm², and 7 for 800 s/mm².

#### 5.2.4. Multifrequency MRE

For multifrequency MRE, mechanical vibrations with frequencies of 300, 400, and 500 Hz were generated by two coupled piezo actuators (APA 200, Cedrat Technologies, France). The vibrations were transferred to the mouse liver through a carbon fiber rod attached to a disc-shaped vibration plate (2.5 cm in diameter), which was placed under the mouse’s abdomen. Shear wave propagation in the liver was recorded at eight dynamics along three motion-encoding directions using a single-shot, spin-echo, echo-planar imaging (SE-EPI) sequence. The frequency and amplitude of the motion-encoding gradient were 299.40 Hz and 34 mT/m, respectively. Twenty slices with an in-plane resolution of 1.0 × 1.0 mm² and 1.2 mm thickness were acquired with TE = 44 ms and TR = 2000 ms. The FoV was 96 × 30 mm².

Figure 5 provides a general overview of the imaging protocol, including a detailed depiction of mouse positioning on the vibration transducer and under the coil, as illustrated in Figure 5(A) and Figure 5(B). Additionally, the vibration actuators are shown in Figure 5(C).

**Figure 5.**
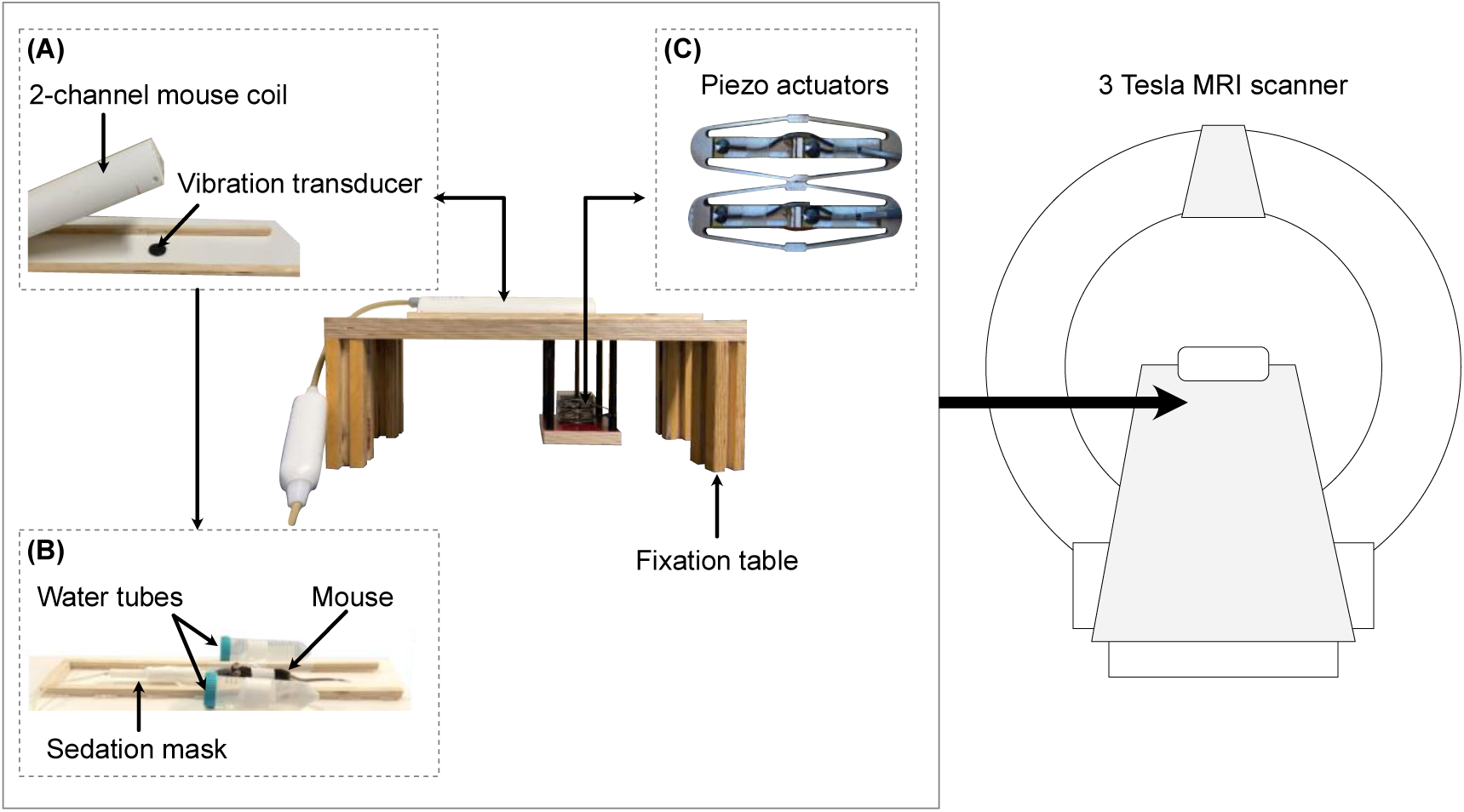
Experimental setup for abdominal imaging of mice in a 3-Tesla scanner. All mice from the five experimental groups were imaged on days 0, 12, 21, 81, and 121 under identical conditions. (A) For imaging, the mouse was placed in prone position beneath a 2-channel coil, with its abdomen resting on a vibration transducer that delivered shear waves to the liver. (B) The mouse was immobilized on a fixation table, connected to an anesthesia source, and surrounded by two water tubes to enhance the imaging signal. (C) The vibration transducer was connected to two coupled piezoelectric actuators. Finally, the entire setup was positioned inside a 3-Tesla MRI scanner.

### 5.3. Image postprocessing

Maps of *HFF* were computed using the IP and opposed-phase OP images acquired with the 2-point Dixon technique:

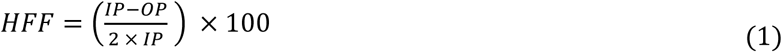

*HFF* maps were represented in grayscale and were scaled from 0 to 100% fat content. *ADC* maps; measured in µm²/s and surrogating water diffusivity; were automatically generated on the scanner workstation using all b-values.

The MRE magnitude image was corrected for breathing artifacts using the statistical parametric mapping (SPM) algorithm [67]. MRE phase images were processed using the wave-number-based multifrequency dual elasto-visco inversion (k-MDEV) technique to generate maps of *SWS* and *PR*, in m/s, as surrogates for tissue stiffness and inverse viscosity, respectively [68]. The MR signal was initially interpolated using bicubic interpolation with a factor of 3 to prevent discretization bias in wavelength estimation [69]. Before phase unwrapping, a low-pass Butterworth filter was applied in the Fourier domain on a zero-padded phase image with an order of 2 and a threshold of 2000 m^-1^. This step was performed to increase resolution in the frequency domain in order to prevent potential Gibbs ringing artifacts. The wavefield resulting after Fourier transform was down-sampled back to the image size before zero-padding, then subjected to a Butterworth high-pass filter (order of 3, threshold of 650 m⁻¹) to suppress compression waves. A directional filter using a cosine-squared function with six directions was applied to the wavefield, which was then used in the final inversion step to produce parameter maps of *SWS* and *PR*, both in m/s. These maps were finally downsampled to their original resolution.

Before registration, all images were cropped to a similar in-plane size (150 × 200 mm²). The MR magnitude images were registered to the corresponding T2w images using Elastix rigid registration [70]. *SWS*, *PR*, and *HFF* maps were then registered to the T2w images using the same transformation matrix. As for the *ADC* maps, the DWI with the lowest b-value was first registered to the T2w image using a sequence of rigid and affine Elastix registration steps. The same transformation matrix from this registration was then applied to the *ADC* maps.

Means of imaging parameters were calculated based on masks that were manually delineated using the T2w anatomical reference. The masks were drawn avoiding the edges and the visible vessels of the liver.

### 5.4. Histology

After imaging, mice were sacrificed via cervical dislocation, and the livers were harvested. The caudate posterior liver lobe was washed with phosphate-buffered saline solution and then snap-frozen for proteomics analysis, while the remaining liver tissue was conserved in a 4% paraformaldehyde solution (Carl Roth, Karlsruhe, Germany) for histopathology.

#### 5.4.1. Semiquantitative histology

Images of H&E-stained liver sections (1 to 2 µm) were used for histological scoring of the progression of MASLD/MASH using the system designed and validated by Kleiner *et al*. [1].

Four histopathological features were evaluated semiquantitatively: steatosis (0-3), lobular inflammation (0-2), hepatocellular ballooning (0-2), and fibrosis (0-4). Upon the diagnosis of MASLD/MASH, the NAS was also calculated, representing the sum of scores for steatosis, lobular inflammation, and ballooning, with a range of 0 to 8 [1].

### 5.4.2. Quantitative Histology

Quantitative histology was performed as a complementary method for quantifying steatosis and lobular inflammation in high-resolution H&E-stained liver sections. Segmented H&E images at 2X magnification with a resolution of 228 × 228 nm² were used.

As a first step for segmentation of steatosis, the high-resolution images were sectioned into approximately 250 tiles, each measuring 5000 × 5000 pixels and encompassing an area of 1140 x 1140 µm² to enhance precision. This was achieved by using the OpenSlide library (version 1.4.1, https://github.com/openslide/openslide-python). For each tile, gamma correction was performed to improve image contrast using the skimage.exposure library (https://scikit-image.org/). Secondly, watershed segmentation was applied to each image tile using the OpenCV library (https://opencv.org/), isolating bright regions that served as seed points for subsequent segmentation of fat droplets. Any objects that were incorrectly segmented were removed based on size and aspect ratio thresholds. Finally, the segmented images were analyzed to determine the area (in µm²) and count of fat droplets. The total fat ratio was calculated by summing the areas of the segmented fat droplets and dividing them by the total area of the liver tissue.

To segment infiltrating inflammatory cells, the StarDist segmentation algorithm, which has been trained for the segmentation of cell nuclei in H&E-stained images, was applied (https://github.com/stardist/stardist). Following the application of a preliminary segmentation, the nuclei of hepatocytes were filtered out from the total mask based on their size (diameter ≥ 7 µm), thus isolating inflammatory cells, mainly lymphocytes. For quantification, the number of inflammatory cells per µm² of liver tissue was measured and presented as a cell-infiltrates density parameter.

### 5.5. Proteomics analysis

#### 5.5.1. Liquid chromatography/mass spectrometry (LC-MS) protocol

Snap-frozen liver samples were processed on an automated workstation (Biomek i7, Beckman Coulter, California, U.S.) using the SP3 protocol with one-step reduction and alkylation [71]. The processed liver samples were analyzed with a mass spectrometer (ZenoTOF 7600, SCIEX, Toronto, Canada) coupled to an ultra-performance liquid chromatography system (ACQUITY M-class, Waters, Massachusetts, U.S.).

For chromatographic separation, tissue digests (200 ng) were loaded onto a 300 µm × 150 mm, 1.8 µm column (HSS T3, Waters, Massachusetts, U.S.) at a flowrate of 5 µL/min and a temperature of 35°C. A 19-minute active gradient was applied, increasing from 1% to 40% mobile phase B (0.1% formic acid in acetonitrile) with mobile phase A (0.1% formic acid in water). Mass spectrometry was performed using an MS/MS acquisition method (Zeno SWATH DIA, SCIEX, Toronto, Canada) with 85 variable-size windows and 11-ms accumulation time. Instrument settings were as follows: pressures of ion source gas 1 and 2 of 12 psi and 60 psi, curtain gas of 55 psi, CAD gas of 7 psi, source temperature of 150°C, and spray voltage of 4500 V.

#### 5.5.2. Protein identification and quantification

Raw data were processed using DIA-NN (version 1.8.1, https://github.com/vdemichev/DiaNN) with default parameters and the following adjustment: fragment ion m/z (range 100 to 1800); mass accuracy of 20 ppm for MS1 and MS2; 7^th^ scan window; match-between-runs: enabled; quantification strategy: Robust LC, high precision. A spectral library-free approach was utilized with annotations based on the *Mus musculus* UniProt database (UP000000589). Results were filtered at a 1% false discovery rate at the precursor level. The mass spectrometry proteomics data have been deposited to the ProteomeXchange Consortium via the PRIDE (https://www.ebi.ac.uk/pride/) partner repository with the dataset identifier PXD060718.

#### 5.5.3. Assessment of metabolic capacities

Hepatic metabolic capacities were assessed using the HEPATOKIN1 model [34] in conjunction with a detailed model of hepatic lipid droplet metabolism [72]. The combination of these models encompassed the primary metabolic pathways involved in carbohydrate, lipid, and amino acid metabolism within hepatocytes, as shown by Berndt *et al.* [34]. The electrophysiological processes at the inner mitochondrial membrane, which describe the generation and utilization of the proton motive force, were modelled using kinetic equations based on the Goldman-Hodgkin-Katz framework [73]. The incorporation of hormone-dependent regulation of liver metabolism through reversible enzyme phosphorylation was achieved using a phenomenological transfer function, as detailed by Bulik *et al.* [36]. Proteomic profiles were employed to generate individual model instantiations following the methodology outlined by Berndt *et al.* [34, 35]. Subsequently, the metabolic physiological functions were defined as maximal fluxes obtained under pre-defined physiological conditions, specifically a plasma glucose concentration range of 3-12 mM. Interdependencies among plasma glucose, plasma hormone, and plasma fatty acid (FA) concentrations were accounted for using experimentally determined transfer functions [36].

#### 5.5.4. Evaluation of protein subsets for inflammation, antioxidant capacity, fibrosis, and cell adhesion

Total LC-MS-based protein profiling identified approximately 6400 protein markers. Based on literature analysis, we confounded in our results four specific subsets of 10 protein markers related to hepatic inflammatory activity, 11 to antioxidant activity, 9 to fibrosis, and 7 to cell adhesion. A quantitative subset ratio was calculated for each mouse at all time points by normalizing the total protein count in the corresponding subset to the sum of all protein counts identified in the liver sample. All markers and their corresponding literature sources are summarized in Table S2 in the supplemental material.

### 5.6. Statistical analysis

This study was designed as a pilot study, for which no a priori effect sizes were available. The sample size per time point was limited to n ≤ 10, which necessitated the use of nonparametric tests for all statistical analyses. For each time point of the disease and each assessed parameter, mean values and standard deviations were calculated. Differences between time points were statistically analyzed using the Kruskal-Wallis test, followed by post hoc pairwise comparisons using the Wilcoxon-Mann-Whitney test. To illustrate the detectable effects given the available sample sizes, effect sizes for the Wilcoxon-Mann-Whitney rank-sum test at a two-sided significance level of 0.05 were calculated. With group sample sizes of 10, 9, and 8, the corresponding effect sizes providing 80% power were 0.862, 0.881, and 0.904, respectively. Spearman correlation was used to assess associations between imaging parameters and histological scores, quantitative histology results, proteomics-based fractions, and maximum metabolic capacities. *p* values ≤ 0.05 were considered statistically significant. All statistical analyses were performed using RStudio (Version 2023.06.0, RStudio, PBC, Boston, MA).

## Abbreviations

ACAC: Acetoacetate
*ADC*: Apparent diffusion coefficient
BHB: Beta-hydroybutyrate
CDAHFD: L-amino acid-defined · high-fat diet
DWI: Diffusion weighted imaging
FA: Fatty acid
k-MDEV: Wave-number-based multifrequency dual elasto-visco inversion
MRE: Magnetic resonance elastography
*PR*: Penetration rate
*SWS*: Shear wave speed
TAG: Triaglycerol
VLDL: very-low-density lipoprotein

## Transcript profiling

The mass spectrometry proteomics data are available on the ProteomeXchange Consortium via the PRIDE (https://www.ebi.ac.uk/pride/) partner repository with the dataset identifier PXD060718.

## Author contributions

Yasmine Safraou: Conceptualization, Investigation, Methodology, Project administration, Data Curation, Validation, Visualization, Formal analysis, Writing – original draft.

Kristin Susan Spirgath: Conceptualization, Investigation, Methodology, Project administration, Data Curation, Writing – original draft.

Biru Huang: Investigation, Methodology, Project administration, Data Curation, Writing – original draft.

Christian Bayerl: Conceptualization, Methodology, Project administration, Resources, Writing – original draft.

Karolina Krehl: Conceptualization, Methodology, Project administration, Writing – original draft.

Anja A. Kühl: Funding acquisition, Investigation, Methodology, Resources, Data curation, Writing – original draft.

Tom Meyer: Methodology, Software, Writing - original draft.

Mehrgan Shahryari: Investigation, Methodology, Data Curation, Validation, Writing – original draft.

Pedro Dantas de Moraes: Data Curation, Writing – original draft.

Jakob Jordan: Methodology, Software, Writing - original draft.

Noah Jaitner: Methodology, Software, Writing - original draft.

Dominik Geisel: Conceptualization, Funding acquisition, Project administration, Resources, Supervision, Writing – original draft.

Jörg Schnorr: Funding acquisition, Project administration, Resources, Writing – original draft.

Nicola Stolzenburg: Conceptualization, Funding acquisition, Project administration, Resources, Supervision, Writing – original draft.

Michael Mülleder: Project administration, Data Curation, Writing – original draft.

Kathrin Textoris-Taube: Project administration, Data Curation, Writing – original draft.

Iwona Wallach: Project administration, Data Curation, Validation, Writing – original draft.

Nikolaus Berndt: Project administration, Conceptualization, Investigation, Methodology, Data Curation, Validation, Formal analysis, Writing – original draft.

Heiko Tzschätzsch: Conceptualization, Methodology, Software, Project administration, Resources, Supervision, Writing – original draft.

Rebecca G. Wells: Methodology, Project administration, Resources, Supervision, Writing – original draft.

Jürgen Braun: Funding acquisition, Conceptualization, Data curation, Investigation, Methodology, Project administration, Validation, Supervision, Writing – original draft.

Patrick Asbach: Funding acquisition, Conceptualization, Data curation, Investigation, Methodology, Project administration, Validation, Supervision, Writing – original draft.

Ingolf Sack: Funding acquisition, Conceptualization, Data curation, Investigation, Methodology, Project administration, Validation, Supervision, Writing – original draft.

Jing Guo: Funding acquisition, Conceptualization, Data curation, Investigation, Methodology, Project administration, Validation, Supervision, Writing – original draft.

## Data transparency statement

Data, analytic methods, and study materials will be made available to other researchers via dedicated online repositories upon request.

## Ethics approval statement

All animal procedures were conducted in accordance with national and institutional regulations and received formal approval from the local authority (Landesamt für Gesundheit und Soziales Berlin, approval number G 0207/20).

## SAGER statement

This study was conducted exclusively in male animals to ensure consistency with previous literature employing similar experimental conditions. While this approach facilitates cross-study comparisons, it may limit the extrapolation to our findings for female animals. Future investigations will evaluate potential sex-based differences.

## Synopsis

Liver viscosity, measured in vivo by magnetic resonance elastography, is a sensitive marker of metabolic dysfunction-associated steatohepatitis (MASH), decreasing before fibrosis onset. In a dietary mouse model of MASH, reduced viscosity was linked to increased inflammation and cell adhesion, and a metabolic shift towards ketogenesis.

## Acknowledgements

We extend our gratitude to Kerstin Rubarth, Ph.D., from the Institute of Biometry and Clinical Epidemiology, Charité - Universitätsmedizin Berlin, for her guidance and assistance with the statistical data analysis.

