## Supplemental Material for "Liver Viscosity Decreases Before the Onset of Fibrosis in Metabolic Dysfunction-Associated Steatohepatitis (MASH)"

[2.] Charité – Universitätsmedizin Berlin, corporate member of Freie Universität Berlin and Humboldt-Universität zu Berlin, Department of Radiology, Chariteplatz 1, 10117 Berlin, Germany

[3.] Charité – Universitätsmedizin Berlin, corporate member of Freie Universität Berlin and Humboldt-Universität zu Berlin, Institute of Medical Informatics, Chariteplatz 1, 10117 Berlin, Germany

[4.] Charité – Universitätsmedizin Berlin, corporate member of Freie Universität Berlin and Humboldt-Universität zu Berlin, Department of Radiology, Hindenburgdamm 30, 12203 Berlin, Germany

[5.] Department of Veterinary Medicine, Institute of Animal Welfare, Animal Behavior and Laboratory Animal Science, Freie Universität Berlin

[6.] Charité – Universitätsmedizin Berlin, corporate member of Freie Universität Berlin and Humboldt-Universität zu Berlin, iPATH.Berlin Core Unit, Hindenburgdamm 30, 12203 Berlin, Germany

[7.] Charité - Universitätsmedizin Berlin, corporate member of Freie Universität Berlin and Humboldt-Universität zu Berlin, Core Facility High Throughput Mass Spectrometry, Berlin, Germany

[8.] German Institute of Human Nutrition Potsdam-Rehbruecke (DIfE), Department of Molecular Toxicology, 14558 Nuthetal, Germany

**Corresponding author:**

Prof. Dr. rer. nat. Jing Guo

### 1. Metabolic Profiling

#### 1.1. Model description

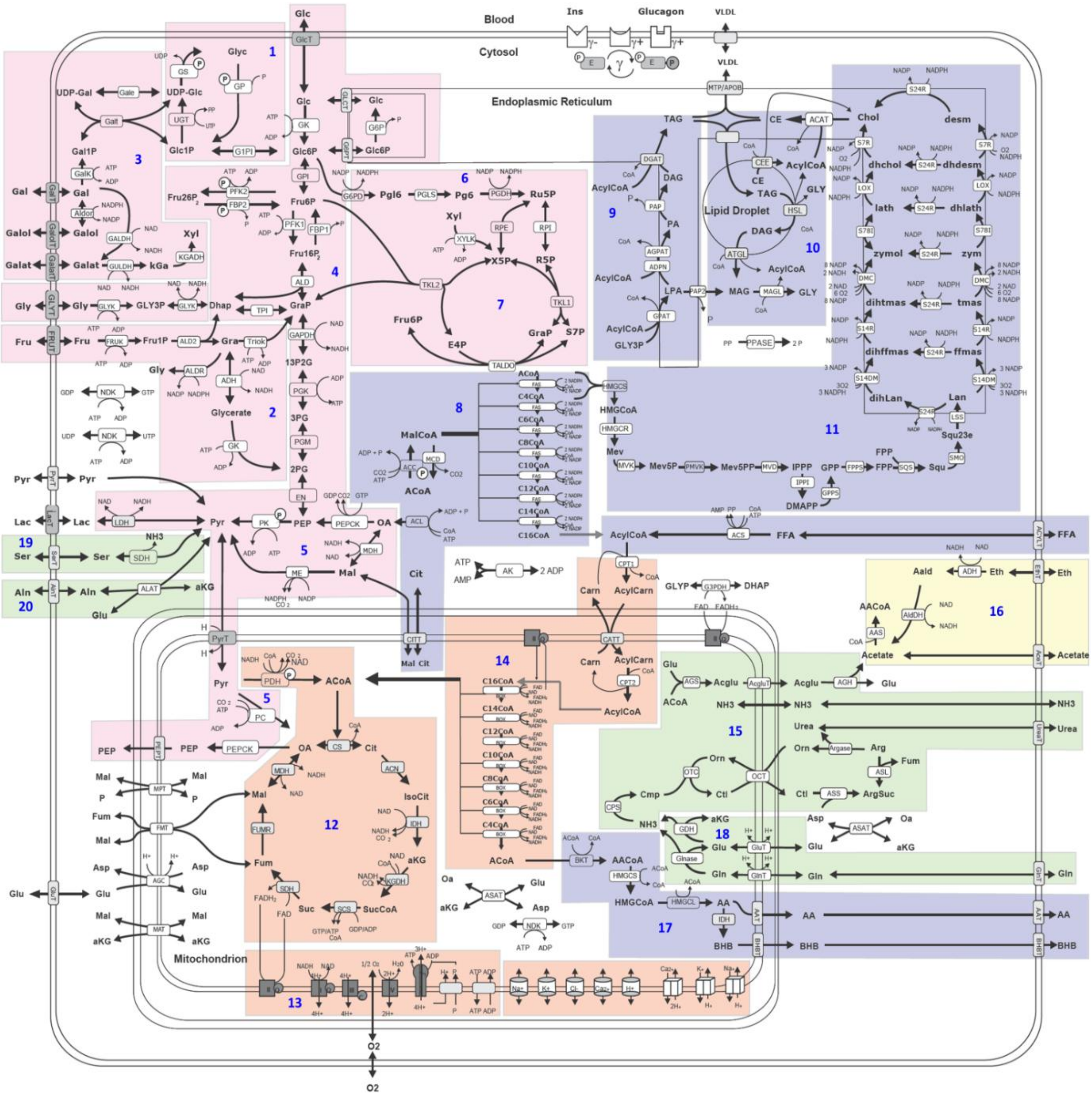


**Figure S1.** Reaction scheme of the metabolic sub-model: Reactions and transport processes between compartments are symbolized by arrows. Single pathways are emphasized by a color-shaded background for the following: (1) glycogen metabolism, (2) fructose metabolism, (3) galactose metabolism, (4) glycolysis, (5) gluconeogenesis, (6) oxidative pentose phosphate pathway, (7) non-oxidative pentose phosphate pathway, (8) fatty acid synthesis, (9) triglyceride synthesis, (10) synthesis and degradation of lipid droplet (LD) synthesis of VLDL lipoprotein, (11) cholesterol synthesis, (12) tricarbonic acid (TCA) cycle, (13) respiratory chain & oxidative phosphorylation, (14)

ß-oxidation of fatty acids, (15) urea cycle, (16) ethanol metabolism, (17) ketone body synthesis, (18) ammonia formation (GlN), (19) serin utilization, and (20) alanine utilization. Small cylinders and cubes symbolize ion channels and ion transporters. Double arrows signify reversible reactions, which may proceed in both directions according to the value of the thermodynamic equilibrium constant and the cellular concentrations of their reactants. Reactions are labeled by the short names of the catalyzing enzyme or membrane transporter given in the small boxes attached to the reactions arrow. Metabolites are denoted by their short names. Full names, kinetic rate laws of reaction rates, a comparison of experimentally determined and calculated metabolic functions, and cellular metabolite concentrations are outlined in [1, 2].

*Metabolic pathways*

The metabolic part of the kinetic model is comprised of the major cellular metabolic pathways, including cellular carbohydrate, lipid, and amino acid metabolism (see Figure S1). The model also contains key electrophysiological processes at the inner mitochondrial membrane, including the membrane transport of various ions, the mitochondrial membrane potential, and the generation and utilization of the proton motive force. The time-dependent variations of the model variables, which equal the concentration of metabolites and ions, are governed by first-order differential equations. Time variations of small ions were modeled by kinetic equations of the Goldman-Hodgkin-Katz type [3]. Numerical values for kinetic parameters of the enzymatic rate laws were taken from reported kinetic studies of the isolated enzyme. Maximal enzyme activities ($V_{max}$ values) were estimated based on a healthy liver’s functional characteristics and metabolite concentrations [1].

*Hormonal signaling*

Hormone-dependent regulation of metabolism by reversible enzyme phosphorylation was accounted for using a phenomenological model of insulin, glucagon-dependent changes of plasma glucose, and changes in the phosphorylation state of interconvertible enzymes as described in [4].

#### 1.2. Individual model parametrization

The establishment of disease-specific metabolic models were based on HEPATOKIN1, a molecularly resolved, physiology-based kinetic model of central metabolism, comprising of the main pathways of energy, carbohydrate, lipid, and amino acid metabolism [1], as well as a molecularly resolved model of lipid droplet metabolism that describes the uptake, storage, and release of triacylglycerol in lipid droplets [2] (see Fig 1).

Individual instantiations of the model were obtained using the protein intensity profiles delivered by quantitative shotgun proteomics to scale the maximal activities of enzymes and transporters. This exploits the fact that the maximal activity of an enzyme is proportional to the abundance of the enzyme protein, according to the relation:

$$v_{max}^{sample} = v_{mx}^{mean\ control} \frac{E^{sample}}{E^{mean\ control}}$$

The maximal activities $v_{mx}^{mean\ control}$ for the control tissue were obtained from [1], $E^{mean\ control}$ denoting the mean protein abundance in the control group, and $E^{sample}$ denoting the protein abundance of enzyme E in the sample.

#### 1.3 Assessment of metabolic capacities: Maximum capacities

The model was used to calculate various metabolic capacities, defined by the magnitude of flux changes in response to changes in the concentration of a distinct plasma metabolite.

**Table S1.** Metabolic capacities and corresponding varied plasma metabolites

| Metabolic Function | Varied Metabolite | Metabolic Function | Varied Metabolite |
|---|---|---|---|
| Fatty acid uptake | Fatty acids | Gluconeogenesis | Lactate |
| Ketone body production | Fatty acids | Glycerol uptake | Glycerol |
| VLDL secretion | Fatty acids | Fructose uptake | Fructose |
| Triglyceride synthesis | Fatty acids | Galactose uptake | Galactose |

| | | | |
|---|---|---|---|
| TAG storage | Fatty acids | Glycogen turnover | Glucose (fasting/refeeding) |
| Beta-oxidation | Fatty acids | Ammonia uptake | Ammonia |
| Ethanol uptake | Ethanol | Urea synthesis | Ammonia |

The varied plasma metabolites, and the monitored corresponding metabolic functions, are listed in Table S1. The non-varied plasma metabolites were kept constant. The external conditions for all simulations are given in [1]. Simulations were run until a steady state was reached.

##### 1.4. Metabolic functions at standardized physiological conditions

The metabolic function was assessed under physiological conditions that corresponded to the mean diurnal standardized plasma profile, meaning that the plasma metabolites were set to the mean value observed throughout a day. The metabolic functions were assessed after the cell reached equilibrium with its surroundings; therefore, the simulation was run until a metabolic steady state was reached.

##### 1.5. Assessment of Metabolic Function in the Range of Physiological Conditions

Metabolic functions were assessed under physiological conditions, which is when plasma metabolite concentrations are not independent of each other. Under physiological conditions, glucose stimulates insulin release from beta cells, concomitants reduce glucagon release from alpha cells in the pancreas, and both hormones control the release of fatty acids from adipose tissue. The interdependence between plasma glucose, plasma hormone, and plasma fatty acid concentrations was considered using the sigmoid Hill-type function, which experimentally determines the glucose-insulin, glucose-glucagon, and glucose-fatty acid relations [1, 2, 4]. Variations of plasma glucose levels were performed under various physiological conditions, ranging from prolonged fasting (3 mM) to excessive nutrient uptake (12 mM plasma glucose). The resulting stationary steady-state metabolic profiles were recorded.

##### 1.6. Diurnal changes in metabolism

The model was used to monitor changes in metabolism in response to time-dependent variations of metabolite and hormone concentrations in the blood plasma profile. Model input was a generic diurnal plasma profile, comprising carbohydrates, fatty acids, amino acids, and hormones. The simulation output was the metabolic state of the sample at any time point of the 24h cycle, resulting in the concentration values of all internal metabolites, internal fluxes, and exchange fluxes.

##### 1.7. Significantly Regulated Functional Proteins

Significantly regulated functional proteins were determined by a linear regression model assessing the association between maximal metabolic capacities and the abundance of proteins belonging to the respective pathway.

#### 2. Statistical Methods

Group values were checked for normality by one-sample Kolmogorov-Smirnov test. Significant differences between groups were assessed by two-sided t-test for normal distributed group values, otherwise, Wilcoxon signed-ranked test was used. Differences between groups are indicated by colored crossbars (yellow: $p < 0.25$; orange: $p < 0.1$; black: $p < 0.05$; red: $p < 0.01$).

#### 3. Significant maximum metabolic functions

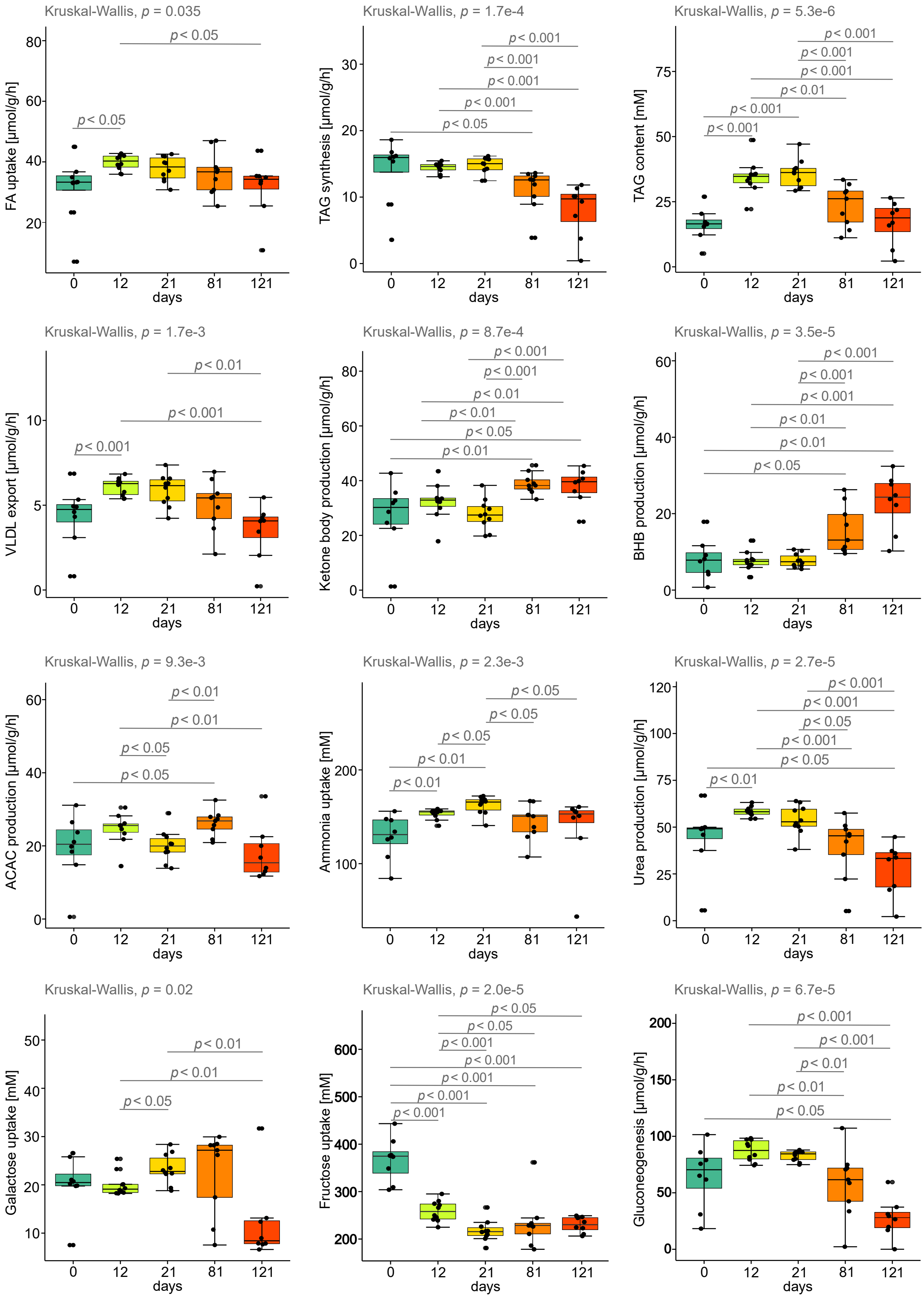

**Figure S1.** Boxplots of evolution of metabolic functions with significant changes throughout the course of disease progression (0, 12, 21, 81, and 121 days). Differences between time points were statistically analyzed using the Kruskal-Wallis test, and post hoc pairwise comparisons were performed using the Wilcoxon-Mann-Whitney test. Lower and upper quartiles and medians are shown. FA stands for fatty acid, TAG for triacylglycerol, VLDL for very low-density lipoprotein, BHB for beta-hydroxybutyrate, and ACAC for acetoacetate.

#### 4. Protein subset markers to evaluate inflammation, antioxidation, fibrosis and cell adhesion

**Table S2.** Representative protein markers linked to key biological processes in Metabolic dysfunction-associated fatty liver disease/metabolic dysfunction-associated steatohepatitis (MAFLD/MASH), including inflammation, antioxidation, cell adhesion, and fibrosis. The markers have been selected from a variety of proteomic studies and literature.

| Subclass | References | Protein markers |
|---|---|---|
| Inflammation | [5] | Csf1r |
| | | TRIL |
| | | Cd163 |
| | | Cd5l |
| | | Chil4 |
| | | Trem2 |
| | | Chil3 |
| | [6] | Tnfaip2 |
| | | Tnfaip8 |
| | | Tnfaipl2 |
| | | Il6st |
| Antioxidation | [7] | Prdx1 |
| | | Prdx4 |
| | | Prdx6 |
| | | Ephx2 |
| | | Gpx1 |
| | | Gpx3 |
| | | Gpx4 |
| | | Dhdh |
| | | Gstp1 |
| | | Haao |
| | | Sod |
| Fibrosis | [8] | Col12a1 |
| | | Col14a1 |
| | | Col15a1 |
| | | Col18a1 |
| | | Col1a1 |

| | | |
|---|---|---|
| | | Col1a2 |
| | | Col3a1 |
| | | Col4a1 |
| | | Col4a2 |
| Cell adhesion | [5] | Col6a1 |
| | | Lgals3bp |
| | | Vwf |
| | | Itga1 |
| | | Itgb1 |
| | | Igfbp7 |
| | | Igfals |

#### 5. Alternative biomechanical parameters

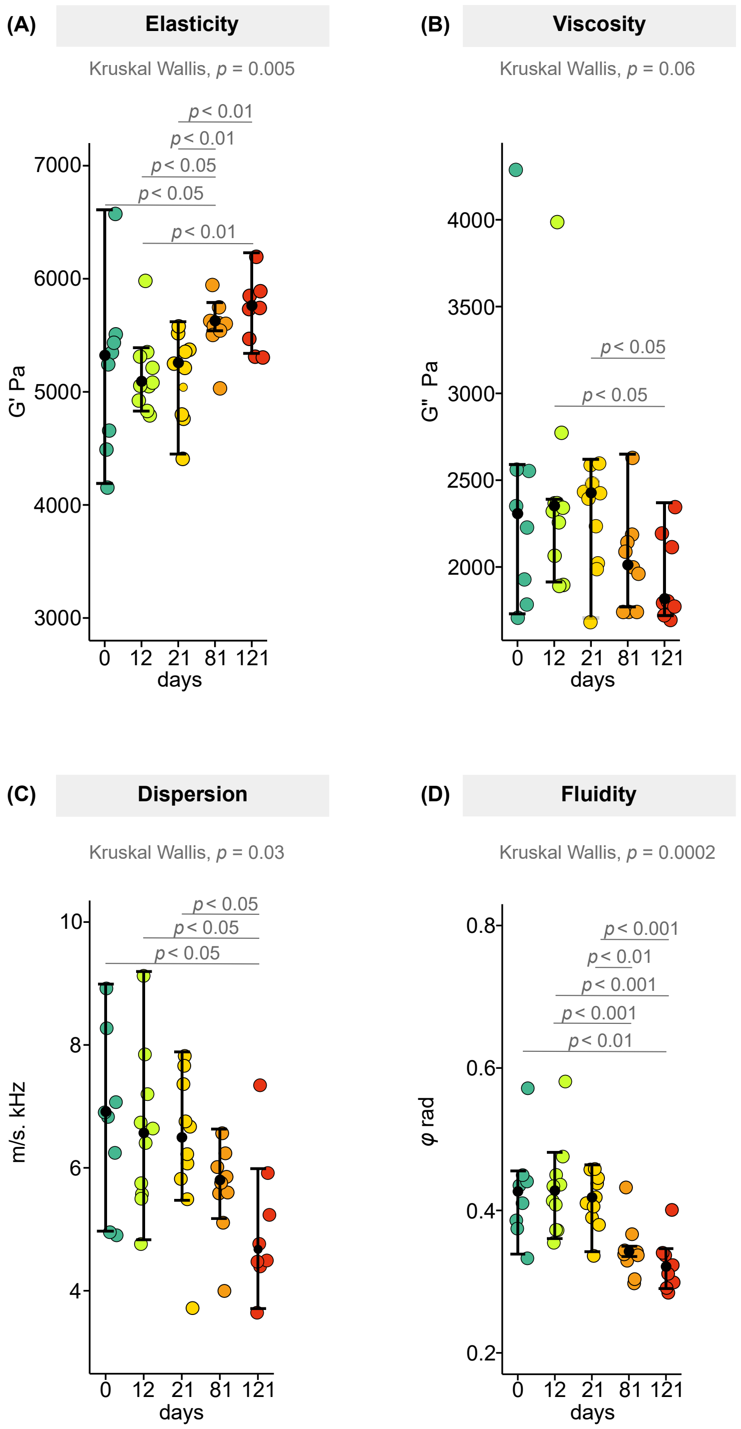


**S4.** Swarm plots showing the progression of alternative mechanical parameters at experimental intervals (0, 12, 21, 81, and 121 days). These parameters include the storage modulus $G'$ (A) , a stiffness surrogate, the loss modulus $G''$ (B), the shear wave speed dispersion (C), and the loss angle

$\varphi$ (D), which surrogate tissue viscosity. Statistical analyses were conducted using the Kruskal-Wallis test, with Wilcoxon Mann-Whitney post-hoc pairwise comparisons. Plots display the median, lower, and upper quartiles for each parameter across all time points.
